# ER shaping proteins regulate mitochondrial fission, outer membrane permeabilization and apoptosis

**DOI:** 10.1101/340448

**Authors:** Mateus Milani, Gerald M Cohen, Shankar Varadarajan

**Affiliations:** Departments of Molecular and Clinical Cancer Medicine, Institute of Translational Medicine, University of Liverpool, Liverpool, Ashton Street, Liverpool, L69 3GE, UK.; Molecular and Clinical Pharmacology, Institute of Translational Medicine, University of Liverpool, Liverpool, Ashton Street, Liverpool, L69 3GE, UK.

**Author notes:** To whom correspondence should be addressed, Shankar Varadarajan, Department of Molecular and Clinical Cancer Medicine, Institute of Translational Medicine, University of Liverpool, 232, Sherrington Building, Ashton Street, Liverpool, L69 3GE, UK.

**Keywords:** Apoptosis, BH3 mimetics, DRP-1, ER shaping proteins, Mitochondrial fission

## Abstract

The mitochondrial fission machinery, comprising a dynamin-related GTPase, DRP-1, is crucial for the regulation of mitochondrial membrane dynamics. Recent reports suggest that the tubular architecture of the endoplasmic reticulum (ER) marks the constriction sites on the mitochondria to facilitate DRP-1-mediated mitochondrial fission. However, the role of several ER shaping proteins that maintain the elaborate network of tubes and sheets in mitochondrial constriction and fission is not yet known. In this report, we demonstrate that modulation of the expression levels of key ER shaping proteins, namely Reticulon1 (RTN-1), Reticulon 4 (RTN-4), Lunapark-1 (LNP-1) and CLIMP-63, markedly decreased the extent of mitochondrial fission mediated by BH3 mimetics, despite no detectable changes in DRP-1 recruitment to the mitochondria. Furthermore, modulation of ER shaping proteins significantly decreased other hallmarks of apoptosis, such as mitochondrial outer membrane permeabilization, caspase activation and phosphatidylserine externalization, and functioned independently of mitochondrial cristae remodeling, thus demonstrating a requirement of ER shaping proteins and ER structural integrity for the efficient execution of the instrinsic apoptotic pathway.

## Introduction

Mitochondria are highly dynamic organelles, which depend on the activities of several membrane-bound fusion and fission GTPases, such as mitofusins, OPA1 and DRP-1, to maintain their structure [1]. Defects in mitochondrial fusion and fission events have been implicated in a range of pathophysiological conditions including poor brain development, optic atrophy, cardiomyopathy and neurodegenerative diseases [2,3]. While the fusion GTPases, such as mitofusins and OPA1, reside on mitochondrial membranes, the fission GTPase, DRP-1 is normally translocated from the cytosol to the outer mitochondrial membrane (OMM) to facilitate mitochondrial fission [4]. The recruitment of DRP-1 to mitochondria has been shown to occur following the marking of mitochondrial constriction sites by the endoplasmic reticulum (ER) [5]. Moreover, several other proteins, such as INF2, Spire1C, actin, Septin 2, Dynamin-2, have been implicated in these events to aid in mitochondrial fission [6–11].

Maintenance of mitochondrial structure is critical for several processes, ranging from metabolism and bioenergetics to intracellular signalling and apoptosis. Most cancer chemotherapeutic agents induce cancer cell death by perturbing mitochondria and activating the intrinsic apoptotic pathway. Several cancers express high levels of the anti-apoptotic BCL-2 family of proteins, such as BCL-2, BCL-X_L_ and MCL-1, which can be inhibited by a new class of highly specific and potent drugs called BH3 mimetics [12,13]. Some of these inhibitors, such as ABT-199 (BCL-2 specific), A-1331852 (BCL-X_L_ specific) and A-1210477 and S63845 (MCL-1 specific) are already approved for use in patients or in clinical trials to treat different malignancies [14–18]. BH3 mimetics perturb mitochondrial integrity by causing mitochondrial outer membrane permeabilization (MOMP) and a concomitant release of cytochrome c, thereby resulting in caspase activation and apoptosis [13,19]. Our recent findings have revealed a role for DRP-1 in BH3 mimetic-mediated mitochondrial fission and apoptosis [20].

In this report, we demonstrate that DRP-1 is required for BH3 mimetic-mediated induction of MOMP and apoptosis. Furthermore, we characterize a critical role for several ER shaping proteins in the execution of BH3 mimetic-mediated mitochondrial fission, MOMP and apoptosis, thus revealing new molecular insights into the mechanisms underlying ER membrane dynamics and cancer cell death.

## Results and Discussion

### DRP-1 is critical for MOMP during BH3 mimetic-mediated apoptosis

Our earlier findings demonstrated a requirement for DRP-1 in BH3 mimetic-mediated apoptosis in several cell lines derived from solid tumors [20]. Since most solid tumors depend on both BCL-X_L_ and MCL-1 for survival, neutralization of both these proteins (using a combination of specific BH3 mimetics) is required to induce apoptosis in these cancers. In agreement, exposure of H1299 cells to a combination of A-1331852 and A-1210477, to inhibit BCL-X_L_ and MCL-1, respectively, resulted in a near-complete release of mitochondrial cytochrome *c* (Fig.1A). This was markedly reduced following a genetic knockdown (siRNA) of DRP-1 or a dominant negative DRP-1 mutant (K38A), thus placing DRP-1 upstream of MOMP (Fig. 1A). Surprisingly, the observed retention of cytochrome *c* occurred in mitochondria that were visibly swollen (Fig. 1A). Support for this was provided by the ultrastructural alterations of mitochondria in cells exposed to BH3 mimetics, which included breaks in the OMM (denoted by the yellow arrowhead), mitochondrial matrix swelling and a concomitant loss of cristae (Fig. 1B). Downregulation of DRP-1 resulted in elongated mitochondria in the control cells and exposure of these cells to BH3 mimetics resulted in visibly swollen mitochondria with intact cristae in the absence of OMM breaks (Fig. 1B).

**Figure 1.**
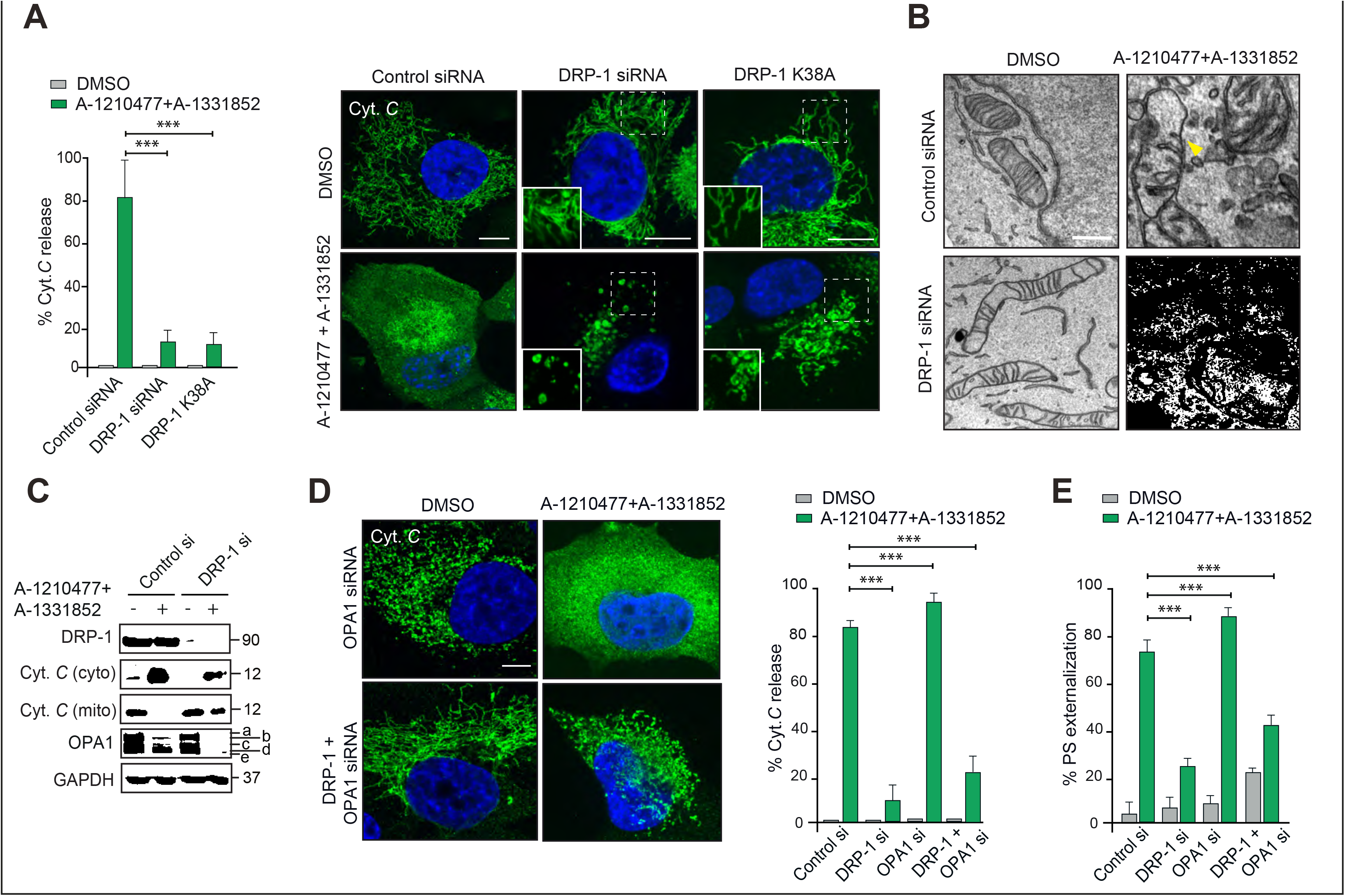
DRP-1 regulates BH3 mimetic-induced mitochondrial fission and apoptosis downstream of mitochondrial cristae remodelling. A H1299 cells, transfected with DRP-1 siRNA or DRP-1 K38A for 72 h, were exposed to Z-VAD.fmk (30 μM) for 0.5 h, followed by a combination of BH3 mimetics, A-1331852 (100 nM) and A-1210477 (10 μM) for 4 h and the extent of cytochrome *c* released from mitochondria was quantified by counting at least 100 cells from three independent experiments. The confocal images in the right panel indicate the inhibitory role of DRP-1 downregulation in BH3 mimetic-mediated release of mitochondrial cytochrome *c*. The boxed regions in the images are enlarged to show mitochondrial structural changes in the indicated cells. B Electron microscopy of H1299 cells, transfected with DRP-1 siRNA for 72 h in the presence or absence of the indicated BH3 mimetics for 2 h. Breaks in the outer mitochondrial membrane are indicated by the yellow arrowhead. Scale bars = 10 nm. C Same as (A), but the extent of cytochrome *c* release as well as OPA-1 processing and the silencing efficiency of DRP-1 siRNA were analyzed by western blotting. D H1299 cells were transfected with the indicated siRNAs for 72 h and treated as (A) and the extent of cytochrome *c* release assessed and quantified. E Same as (D) but the cells were exposed to BH3 mimetics in the absence of Z-VAD.fmk and the extent of apoptosis assessed by PS externalization. Data information: The data in all graphs are presented as Mean ± SEM. ***P⩽0.001 (oneway ANOVA). All Scale bars, unless indicated otherwise correspond to 10 μm.

### DRP-1 acts downstream of cristae remodelling during BH3 mimetic-mediated apoptosis

MOMP is generally accompanied by the release of mitochondrial cytochrome *c* from the inner membrane space to the cytosol, as a consequence of mitochondrial cristae remodelling [21]. Exposure to BH3 mimetics caused a loss of the high molecular weight isoforms of OPA1, characteristic of mitochondrial cristae remodelling [22,23], which was accompanied by an almost complete release of mitochondrial cytochrome *c* into the cytosol (Fig. 1C). Downregulation of DRP-1 markedly diminished the release of mitochondrial cytochrome *c* but failed to prevent OPA1 proteolysis, induced by the combination of BH3 mimetics (Fig. 1C). Downregulation of DRP-1 not only prevented the mitochondrial fission following the loss of fusion in OPA1 siRNA-transfected cells but also inhibited the complete release of cytochrome *c*, following exposure to the BH3 mimetics in the OPA1 downregulated cells (Fig. 1D). In an analogous manner, downregulation of Drp-1 inhibited the extensive apoptosis, assessed by phosphatidylserine (PS) externalization even following downregulation of OPA1 (Fig. 1E). Thus these data indicate that DRP-1 plays a role downstream of cristae remodelling and also upstream of cytochrome *c* release.

### ER shaping proteins regulate BH3 mimetic-mediated mitochondrial fission

Since the ER tubules have been shown to mark the constriction sites on the mitochondria to facilitate DRP-1-mediated fission [5], we speculated whether modulating the expression levels of ER shaping proteins could alter mitochondrial membrane dynamics and function. To test this hypothesis, we genetically silenced the expression levels of some of key ER shaping proteins, namely Reticulon1 (RTN-1), Reticulon4 (RTN-4), Lunapark-1 (LNP-1) and CLIMP-63 in H1299 cells. Of these, RTN-1 and RTN-4 contribute to the curvature and maintenance of ER tubules, whereas CLMP-63 helps to maintain ER sheets [24,25]. LNP-1 is localized to the three-way junctions in the ER and contributes to ER network formation [26,27]. In cells transfected with siRNAs targeting these ER shaping proteins, we wished to assess changes in mitochondrial structure, following inducers of mitochondrial fission, such as A-1210477 (MCL-1 inhibitor) and CCCP (mitochondrial uncoupler). Downregulation of DRP-1 prevented both A-1210477- and CCCP-mediated mitochondrial fission, whereas downregulation of the different ER shaping proteins markedly inhibited A-1210477-but not CCCP-mediated mitochondrial fission (Figs. 2 A-C). This may be due to the different mechanisms by which A-1210477 and CCCP induce mitochondrial fission, with CCCP inducing mitochondrial depolarization and Parkin-dependent mitophagy [28]. How MCL-1 regulates mitochondrial membrane dynamics is largely unknown [29–33]. In H1299 cells, A-1210477 neither induced mitochondrial depolarization nor mitophagy, despite causing marked mitochondrial fission [20]. Furthermore, downregulation of DRP-1 but not ER shaping proteins prevented the loss of fusion, following OPA1 siRNA (Figs. 3 A and B). Taken together, these results suggested that ER shaping proteins, similar to DRP-1, regulate BH3 mimetic-mediated mitochondrial fission but unlike DRP-1, ER shaping proteins are dispensable for mitochondrial fission events induced by CCCP and OPA1 downregulation.

**Figure 2.**
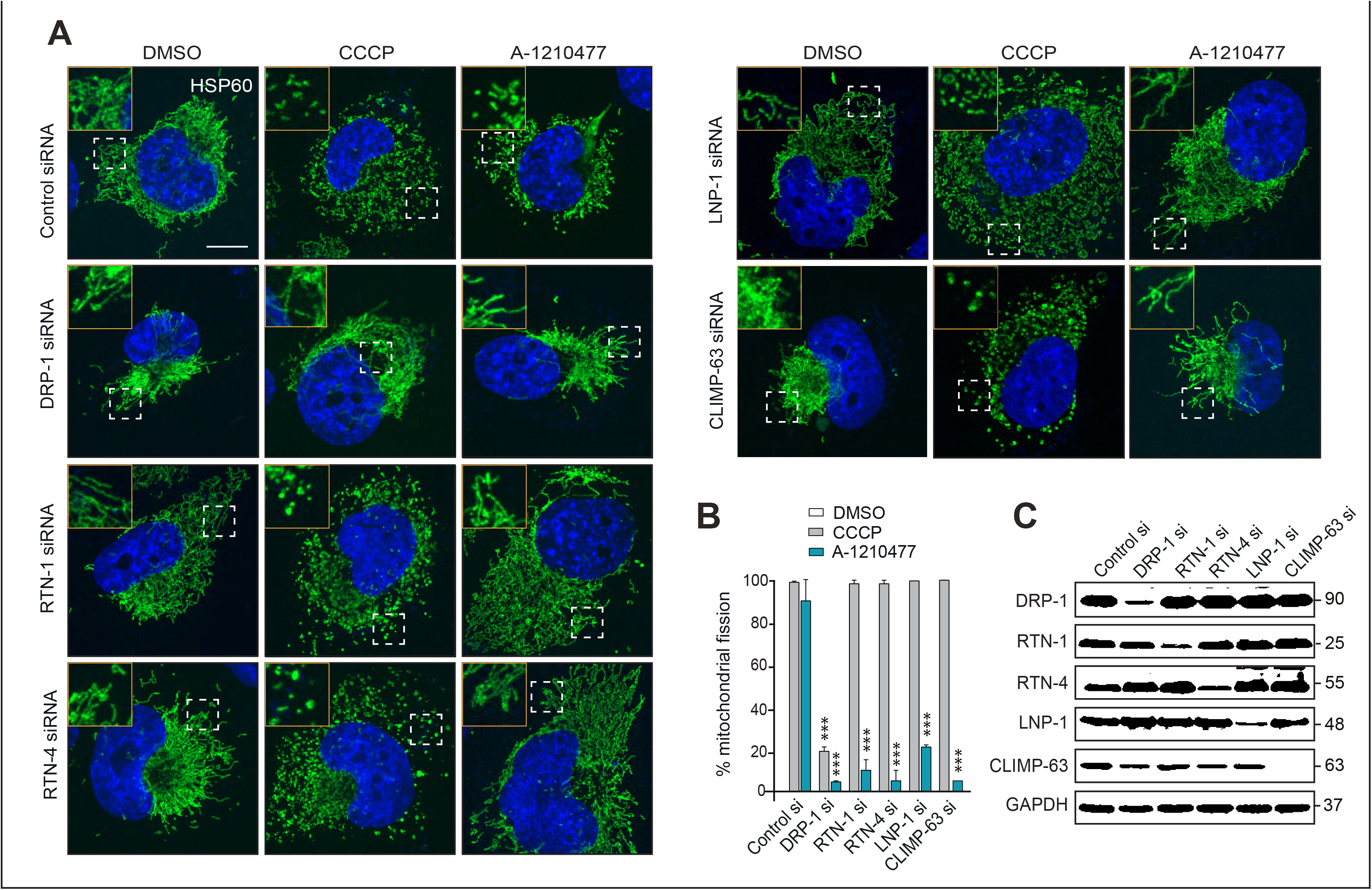
ER shaping proteins are required for A-1210477-but not CCCP-mediated mitochondrial fission. A H1299 cells transfected with the indicated siRNA for 72 h were exposed to DMSO, CCCP (20 μM, 2 h) or A-1210477 (10 μM, 4 h) and the extent of mitochondrial fission assessed by HSP60 staining. The boxed regions in the images are enlarged to show mitochondrial structural changes in the indicated cells. B The extent of fission in these cells was then quantified by counting at least 100 cells from three independent experiments. C Western blots showing the silencing efficiency of the different siRNAs. Data information: The data in the graph are presented as Mean ± SEM. ***P⩽0.001 (one-way ANOVA). Scale bars correspond to 10 μm.

**Figure 3.**
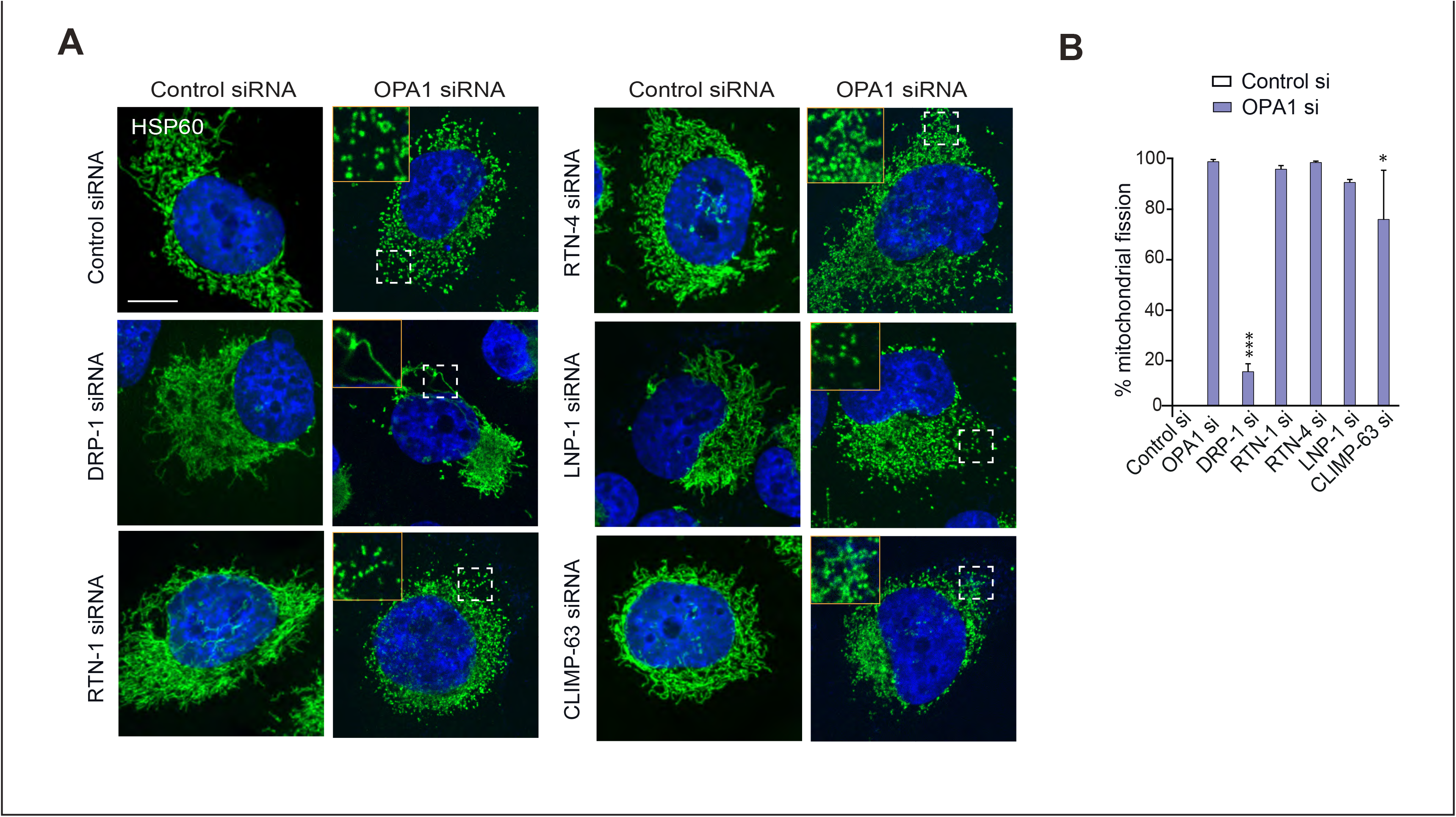
Silencing of DRP-1 but not ER shaping proteins rescues mitochondrial structural changes due to loss of fusion. A H1299 cells were transfected with either control, DRP-1, RTN-1, RTN-4, LNP-1 or CLIMP-63 siRNA alone or in combination with OPA-1 siRNA and assessed for mitochondrial integrity by HSP60 staining. The boxed regions in the images are enlarged to show mitochondrial structural changes in the indicated cells. Scale bars = 10 μm. B The extent of fission in these cells was then quantified by counting at least 100 cells from three independent experiments. Data information: The data in the graph are presented as Mean ± SEM. \**p* ≤ 0.05 and ***P⩽0.001 (one- way ANOVA). Scale bars correspond to 10 μm.

### ER shaping proteins do not affect DRP-1 recruitment to mitochondria

Next, we wished to assess whether the ER shaping proteins regulated mitochondrial fission by facilitating DRP-1 translocation and recruitment to the mitochondrial constriction sites. Surprisingly, downregulation of the ER shaping proteins did not affect DRP-1 recruitment to mitochondrial membranes (Fig. 4). Exposure of control cells to A-1210477 resulted in extensive mitochondrial fission and in these cells DRP-1 was detected proximal to the fissed mitochondrial filaments (Fig. 4). Downregulation of the ER shaping proteins markedly inhibited A-1210477-mediated mitochondrial fission (Figs. 2B and 4), despite DRP-1 being extensively associated with the filamentous mitochondria (Fig. 4). These results implied that recruitment and association of DRP-1 alone to the mitochondrial membrane was insufficient to induce mitochondrial fission and efficient fission required the involvement of ER shaping proteins. However, the absence of ER shaping proteins may alter the phosphorylation status and/or the resident time of DRP1 in the mitochondrial membrane.

**Figure 4.**
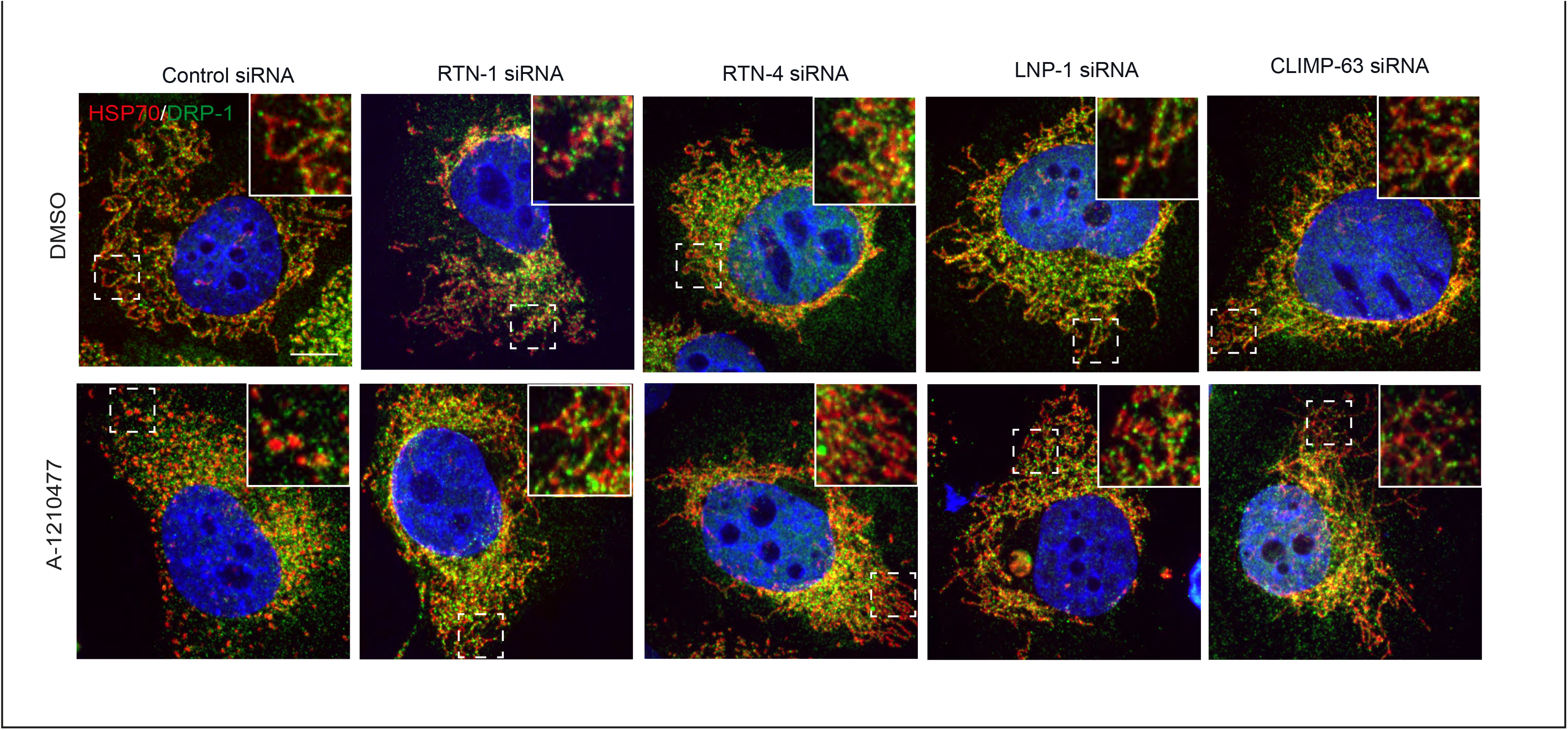
DRP-1 localizes to the mitochondrial fission sites independently of ER shaping proteins. H1299 cells, transfected with the indicated siRNA for 72 h, were exposed to DMSO or A-1210477 (10 μM, 4 h) and immunostained for DRP-1 and HSP70. The boxed regions in the images are enlarged to show DRP-1 localization to the mitochondria in the indicated cells. Data information: Scale bars =10 μm.

### ER shaping proteins prevent BH3 mimetic-mediated MOMP and apoptosis

Since the downregulation of ER shaping proteins antagonized A-1210477-mediated mitochondrial fission similar to that of DRP-1 siRNA (Figs. 2 and 4), we wished to investigate whether silencing the expression of ER shaping proteins could also prevent BH3 mimetic-mediated MOMP and apoptosis. Indeed, downregulation of several ER shaping proteins, such as RTN-1, RTN-4 and CLIMP-63 but not LNP-1, prevented BH3 mimetic-mediated MOMP, characterized by the release of mitochondrial cytochrome *c* (Fig. 5A and B). Since the reticulons and CLIMP-63 maintain ER tubules and sheets along with LNP-1, which forms the three-way junction to maintain the interconnected ER network, the inability of LNP-1 downregulation to prevent BH3 mimetic-mediated MOMP was surprising, and in stark contrast to its potential to prevent BH3 mimetic-mediated mitochondrial fission (Figs. 2 and 5). How or why LNP-1 uncouples BH3 mimetic-mediated mitochondrial fission and MOMP needs further characterization.

**Figure 5.**
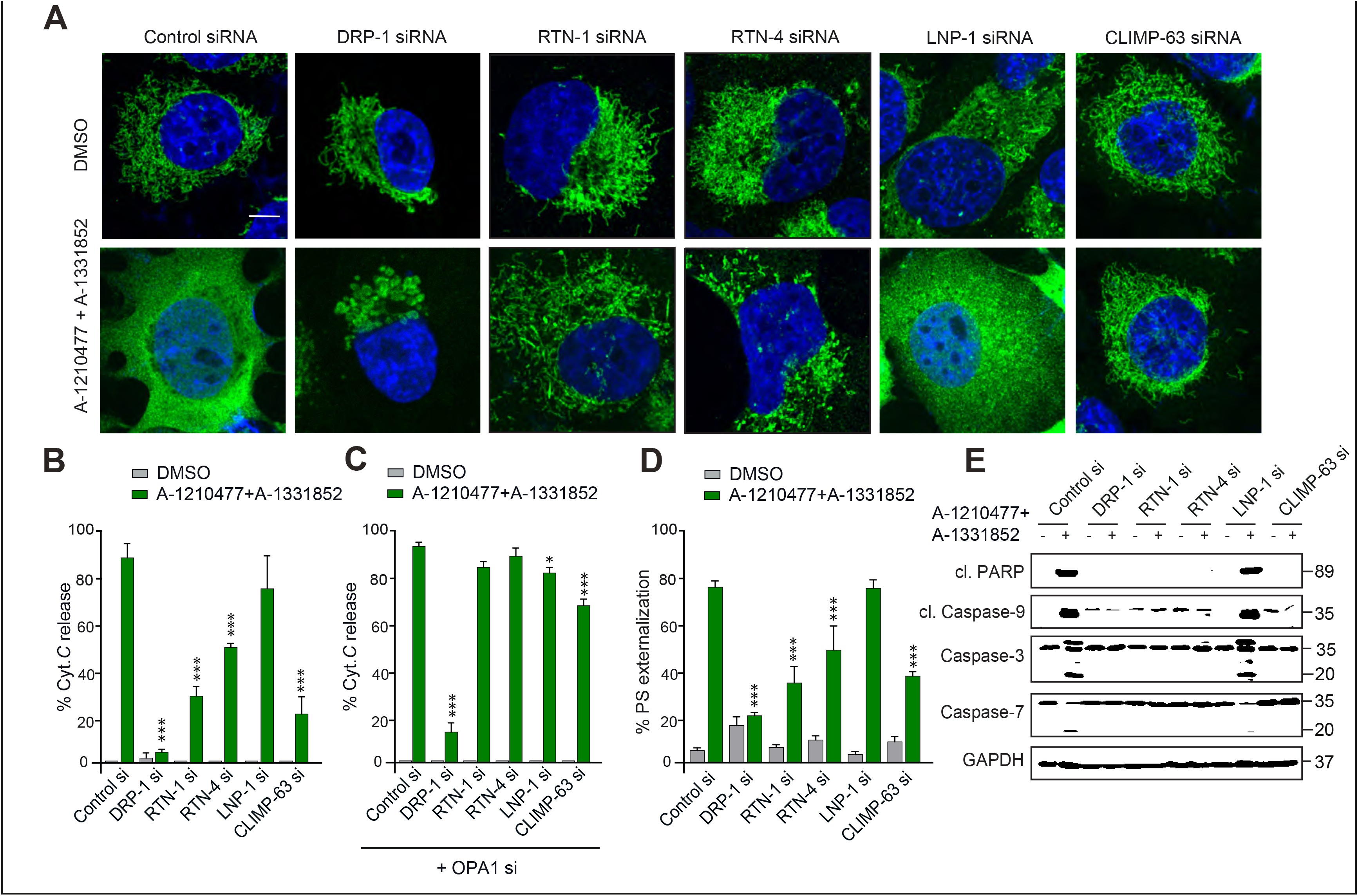
ER shaping proteins antagonize BH3 mimetic-mediated mitochondrial outer membrane permeabilization and apoptosis upstream of cristae remodelling. A H1299 cells were transfected with the indicated siRNAs for 72 h and exposed to Z-VAD.fmk (30 μM) for 0.5 h, followed by a combination of BH3 mimetics, A-1331852 (100 nM) and A-1210477 (10 μM) for 4 h and the extent of cytochrome *c* released from mitochondria was monitored. B At least 100 cells from (A) were quantified for the extent of cytochrome *c* release from three independent experiments. C Same as (A) but the experiments were performed in cells which were also transfected with OPA1 siRNA for 72 h. D and E Same as (A) but the cells were exposed to BH3 mimetics in the absence of Z-VAD.fmk and the extent of apoptosis assessed by (D) PS externalization and (E) western blotting for the activation of caspases and the cleavage of poly(ADP-ribose) polymerase (PARP). Data information: The data in the graph are presented as Mean ± SEM. \**p* ≤ 0.05 and ***P⩽0.001 (one-way ANOVA). Scale bars correspond to 10 μm.

Nevertheless, since DRP-1 facilitated BH3 mimetic-mediated apoptosis downstream of mitochondrial cristae remodeling (Fig. 1E), we wondered whether ER shaping proteins also regulated apoptosis downstream and/or independent of OPA1 proteolysis. Downregulation of ER shaping proteins, unlike DRP-1 siRNA, resulted in a near-complete release of cytochrome c, in cells transfected with OPA1 siRNA (Fig. 5C), suggesting that the ER shaping proteins, in contrast to DRP-1, regulated mitochondrial dynamics upstream of OPA1 proteolysis. This is consistent with the mitochondrial structural changes observed following BH3 mimetics in the different siRNA transfected cells (Fig. 5A). While downregulation of DRP-1 resulted in significant mitochondrial swelling following BH3 mimetics, none of the siRNAs targeting the ER shaping proteins resulted in similar mitochondrial swelling (Fig. 5A), thus supporting the notion that ER shaping proteins, unlike DRP-1, regulated mitochondrial dynamics upstream of IMM swelling. Finally, silencing the expression of most of the ER shaping proteins, except LNP-1 prevented, to varying extents, other hallmarks of apoptosis, such as PS externalization (Fig. 5D) and caspase activation, as assessed by the cleavage of the apical caspae-9 in the intrinsic pathway of apoptosis as well as the effector caspases-3 and −7 (Fig. 5E), following exposure to BH3 mimetics. Taken together, our results demonstrate that BH3 mimetics induced apoptosis in a DRP-1-dependent manner, in which the ER membranes marked the constriction sites on the mitochondria, which in turn facilitated mitochondrial cristae remodeling (OPA1 proteolysis), inner mitochondrial membrane swelling, OMM breaks, cytochrome *c* release, caspase activation and PS externalization, thus highlighting a crucial role for the integrity of the mitochondrial fission machinery and ER membranes in the efficient execution of BH3 mimetic-mediated apoptosis. As BH3 mimetics are currently entering the clinic due to their exquisite specificity and potency to induce apoptosis by antagonizing the anti-apoptotic BCL-2 family members, knowledge of the interplay between the ER and mitochondria dynamics can lead to more effective therapy.

## Materials and Methods

### Cell culture

H1299 cells purchased from ATCC (Middlesex, UK) and authenticated by short tandem repeat (STR) profiling were cultured in RPMI 1640 medium supplemented with 10 % fetal calf serum (FCS) from Life Technologies, Inc, (Paisley, UK) and maintained at 37°C in a humidified atmosphere of 5% CO_2_.

### Reagents

BH3 mimetics, A-1331852 and A-1210477 were kindly provided by AbbVie Inc (IL, USA). CCCP was obtained from Selleck Chemicals Co. (Houston, TX, USA). Antibodies against HSP60, Cytochrome c, OPA1 and DRP-1 from BD Biosciences (San Jose, CA, USA), RTN-1, RTN-4, HSP70 from Abcam (Cambridge, UK), CLIMP-63 from Enzo Lifesciences (Exeter, UK), LNP-1 from Sigma, cleaved PARP, cleaved caspase 9, caspase-3 and caspase-7 from Cell Signaling Technologies (Danvers, MA, USA) and GAPDH from SantaCruz Biotechnologies (Santa Cruz, CA, USA) were used. All other reagents, unless mentioned otherwise, were from Sigma-Aldrich Co. (St. Louis, MO, USA).

### Transfections

For RNA interference, cells were transfected with 10 nM of siRNAs against DRP-1 (SI04274235), RTN-1 (SI04178545), RTN-4 (s32767), CLIMP-63 (SI00347242), LNP-1 (SI00460355) and OPA1 (SI03019429) purchased from either Life Technologies or Qiagen Ltd. (Manchester, UK), using Interferin (Polyplus Transfection Inc, NY), according to the manufacturer’s protocol and processed 72 h after transfection. For transient overexpression, cells were transfected with GFP-DRP-1-DN (provided by Dr. E. Bampton, University of Leicester, UK) using Lipofectamine LTX reagent (Thermo Fisher Scientific, Loughborough, UK) and incubated for 48 h, according to the manufacturer’s instructions.

### Microscopy

For immunofluorescent staining, cells grown on coverslips were fixed with 4 % (w/v) paraformaldehyde, permeabilized with 0.5 % (v/v) Triton X-100 in PBS, followed by incubations with primary antibodies, the appropriate fluorophore-conjugated secondary antibodies, mounted on glass slides and imaged using a 3i Marianas spinning disk confocal microscope, fitted with a Plan-Apochromat 63x/1.4NA Oil Objective, M27 and a Hamamatsu ORCA-Flash4.0 v2 sCMOS Camera (all from Intelligent Imaging Innovations, GmbH, Germany). For transmission electron microscopy, cells were prepared based on the Deerinck and Ellisman SBF SEM protocol [34]. Electron micrographs were recorded using a MegaViewIII CCD camera and AnalySIS software (EMSIS GmbH, Germany) in a FEI Tecnai G2 Spirit BioTWIN electron microscope. Acquired images were processed using ImageJ software (NIH, Bethesda, MD).

### Flow cytometry

Cells were collected, washed with PBS and resuspended in 500 μL of Annexin binding buffer containing 150 mM NaCl, 1 mM MgCl_2_, 5 mM KCl, 2 mM CaCl_2_, 10 mM HEPES, pH 7.4 and Annexin-FITC and incubated for 8 min. This was followed by the addition of 2.5 μg of PI and further incubated for 2 min and apoptosis was assessed by measuring the extent of phosphatidylserine externalization, using an Attune NxT flow cytometer (ThermoFisher Scientific, Paisley, UK).

### Cytochrome *c* release assay

Roughly a million cells were collected, washed twice with PBS at 4 °C and the pellets resuspended in 100 μL of lysis buffer containing 250 mM sucrose, 5 mM MgCl_2_, 10 mM KCl, 1 mM EDTA, 1 mM EGTA, 20 mM HEPES, 0.01 % digitonin, pH 7.4 at 4°C and incubated on ice for 5 min. The cells were then centrifuged at 14,000 g for 3 min at 4 °C and the supernatant collected. The pellet was resuspended in lysis buffer without digitonin and centrifuged again at 14,000 g for 3 min at 4°C. The supernatant was pooled with the previously collected supernatant to form the cytosolic fraction. The remaining pellet was resuspended in lysis buffer without digitonin and sonicated to form the mitochondrial fraction. Western blotting was then carried out in corresponding fractions to assess the extent of cytochrome *c* released from the mitochondria to the cytosol.

### Western blotting

Western blotting was carried out according to standard protocols. Briefly, 50 μg of total protein lysate was subjected to SDS-PAGE electrophoresis. Subsequently proteins were transferred to nitrocellulose membrane and protein bands visualized with ECL reagents (GE Healthcare).

### Statistical Analysis

For time-course studies, a two-way ANOVA was performed and other studies were analyzed for statistical significance with one-way ANOVA and the asterisks depicted correspond to the following *p* values: * *p* ≤ 0.05, ** *p* ≤ 0.005 and *** *p* ≤ 0.001.

## Acknowledgements

We thank AbbVie for providing different BH3 mimetics used in this study. We thank Ms. A.J. Beckett and Prof. I. Prior for assistance with electron microscopy. This work was supported by North West Cancer Research Grant CR1040 (to SV and GMC) and a Science Without Borders, CNPq 233624/2014-7, Ministry of Education, Brazil (to MM). The authors declare no competing financial interests.

## Author Contributions

Mateus Milani performed and analyzed all cellular experiments; Gerald M. Cohen and Shankar Varadarajan wrote the manuscript and supervised all aspects of the study.

